# Pantothenate auxotrophy in *Zymomonas mobilis* ZM4 is due to a lack of aspartate decarboxylase activity

**DOI:** 10.1101/144709

**Authors:** J. R. Gliessman, T. A. Kremer, A. A. Sangani, S. E. Jones-Burrage, J. B. McKinlay

## Abstract

The bacterium *Zymomonas mobilis* naturally produces ethanol at near theoretical maximum yields, making it of interest for industrial ethanol production. *Z. mobilis* requires the vitamin pantothenate for growth. Here we characterized the genetic basis for the *Z. mobilis* pantothenate auxotrophy. We found that this auxotrophy is due to the absence of a single gene, *panD*, encoding aspartate-decarboxylase. Heterologous expression of *Escherichia coli* PanD in *Z. mobilis* or supplementation of the growth medium with the product of PanD activity, β-alanine, eliminated the need for exogenous pantothenate. We also determined that IlvC, an enzyme better known for branched-chain amino acid synthesis, is required for pantothenate synthesis in *Z. mobilis*, as it compensates for the absence of PanE, another pantothenate synthesis pathway enzyme. In addition to contributing to an understanding of the nutritional requirements of *Z. mobilis*, our results have led to the design of a more cost-effective growth medium.

## Introduction

*Zymomonas mobilis* is a bacterium best known as a potential rival to the ethanol-producing yeast *Saccharomyces cerevisiae. Z. mobilis* ferments glucose into ethanol at 97% of the theoretical maximum yield, produces ethanol 3 – 5 times faster than yeast on a per cell basis, and produces less residual biomass than yeast (Jeffries 2005). Unlike yeast, *Z. mobilis* can also use inexpensive N_2_ gas a nitrogen source, raising the possibility to grow *Z. mobilis* on nitrogen-poor cellulosic feedstocks without the need for expensive undefined nitrogen supplements, such as corn steep liquor (Kremer *et al*. 2015). However, these undefined supplements can also satisfy vitamin requirements (Lawford and Rousseau 1997). The vitamin pantothenate (vitamin B5), a precursor to coenzyme-A, is required by all *Zymomonas* isolates characterized to date (Belaich and Senez 1965; De Ley and Swings 1976; Nipkow, Beyeler and Fiechter 1984; Galani, Drainas and Typas 1985; Lawford and Stevnsborg 1986; Cross and Clausen 1993). Elimination of this auxotrophy, combined with utilizing N2 as a nitrogen source, would circumvent the need for nutrient rich supplements altogether for *Z. mobilis* growth on cellulosic feedstocks.

Herein, we describe the genetic basis for panthothenate auxotrophy in the most commonly used *Z. mobilis* research strain, ZM4 (Seo *et al*. 2005; Skerker *et al*. 2013). A comparative genomics analysis of the seven sequenced *Z. mobilis* isolates indicated that the entire pantothenate synthesis pathway (depicted in Fig. 1) is missing in two isolates while the other five strains, including ZM4, are only missing *panD* and *panE* (Fig. 2). We found that β-alanine could support ZM4 growth in place of pantothenate. In support of this observation, expression of a heterologous PanD, the enzyme producing β-alanine from aspartate (Fig. 1), eliminated the pantothenate auxotrophy. We also discovered that the lack of *panE* was inconsequential for pantothenate synthesis as the activity was compensated for by IlvC, an enzyme better known for its role in branched-chain amino acid synthesis. Our results indicate that β-alanine can serve as a less expensive growth supplement in place of pantothenate and that heterologous expression of a single gene, PanD, is sufficient to eliminate the pantothenate auxotrophy.

**Fig 1.**
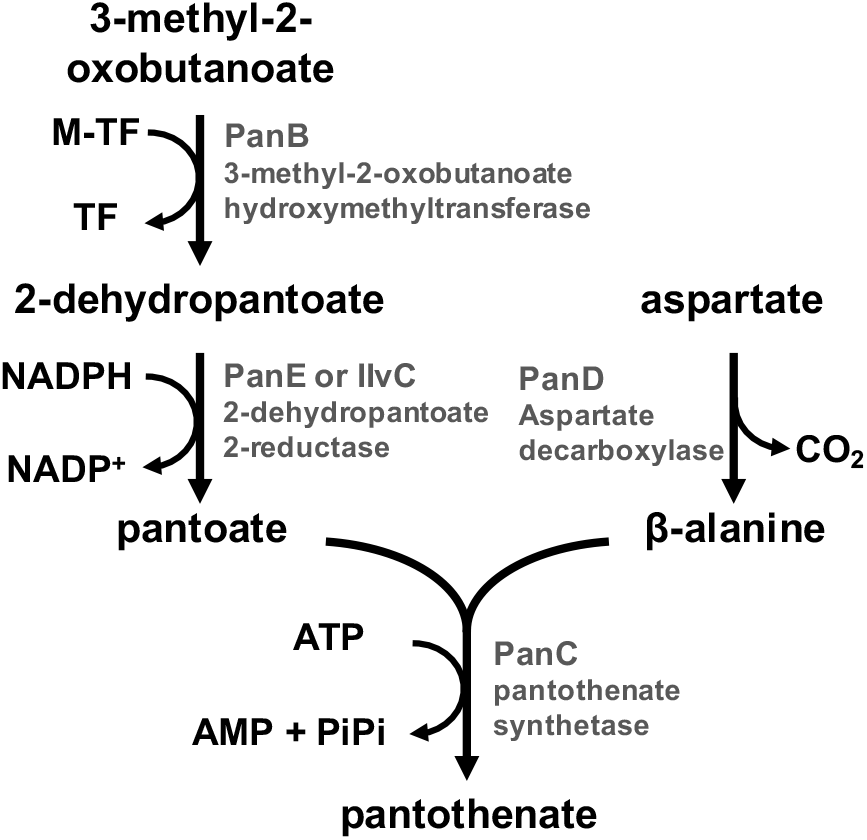
Pantothenate synthesis pathway. 3-Methyl-2-oxobutanoate is derived from valine. M-TF, 5,10-methylene-tetrhydrofolate; TF, tetrahydrofolate.

**Fig 2.**
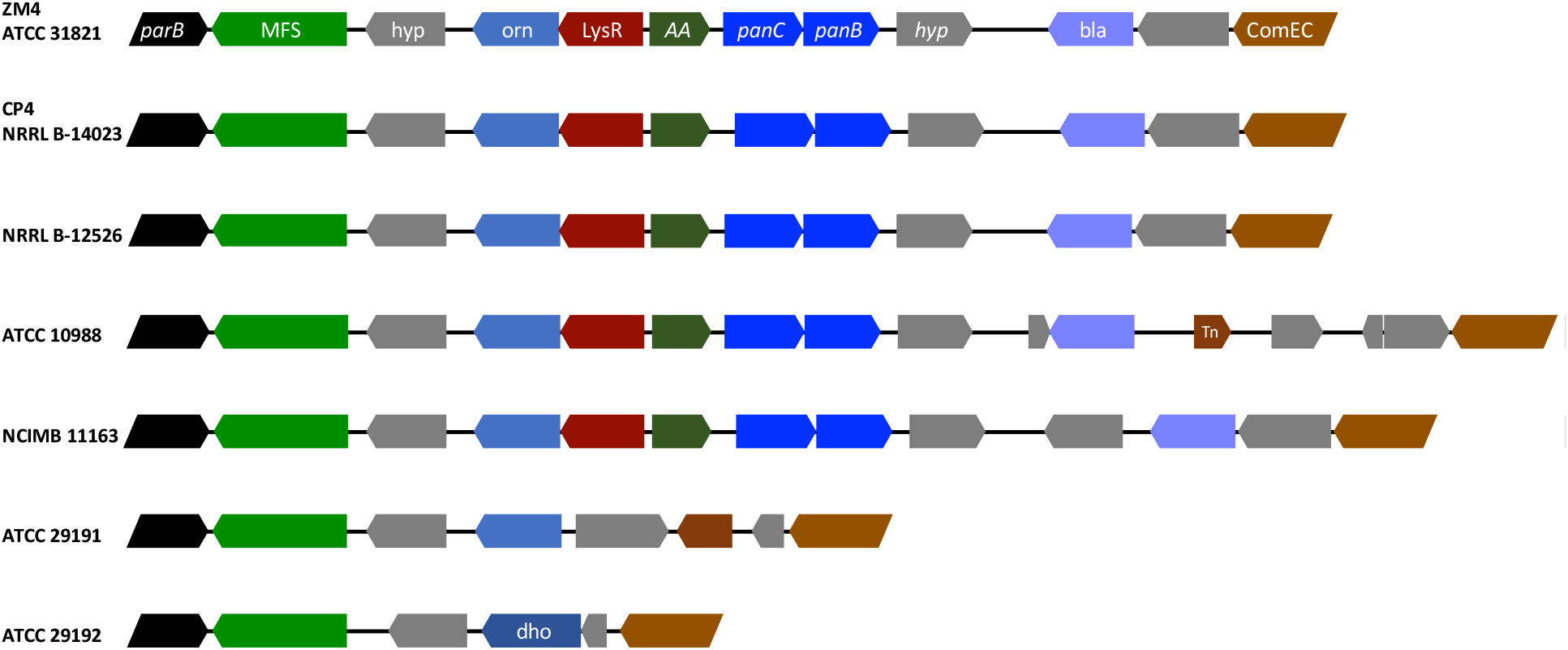
Synteny of *panB* and *panC* genomic regions in the seven sequence *Z. mobilis* strains. Arrow color indicates gene function. Black, *parB*; Light green, major facilitator superfamily transporter; grey, hypothetical protein; Blue, enzymes: orn, ornithine cyclodeaminase; dho, dihydroorotate oxidase; bla, β-lactamase; Red, LysR-family regulator; Dark green, amino acid transporter (AA); light brown, ComEC competence protein; dark brown, transposon (Tn). In ZM4, the locus tags for *panB* and *panC* are ZMO1970 and ZMO1954, respectively. The *panBC* homologs, respectively ZMO1952 and ZMO1971, are located 22.4 kb away.

## Materials and Methods

### Strains and growth conditions

All strains are described in Table 1. *Z. mobilis* ZM4 (ATCC 31821) and the IlvC transposon mutant (ZMO1141::Tn5; UP33_A10) were kindly given to us by J. M. Skerker and A. P. Arkin, UC Berkeley (Skerker *et al*. 2013). For cloning experiments, *Z. mobilis* was grown in aerobic YPG (1% yeast extract, 2% peptone, 2% glucose) or plated on YPG agar (1.5% agar). All growth experiments were conducted in 10 ml of a chemically-defined growth medium (ZYMM) in anaerobic test tubes with shaking at 150 rpm as described (Kremer *et al*. 2015). Where indicated, calcium pantothenate and β-alanine were added at a final concentration of 100 nM each, and branched chain amino acids (isoleucine, leucine, and valine) were added at a final concentration of 0.5 mM each. Media were made anaerobic by bubbling with Ar gas and then sealing tubes with rubber stoppers (Geo-Microbial Technologies, Ochelata, OK) and aluminum crimps. Starter cultures were inoculated with a single colony from YPG agar, and then a 1% inoculum was transferred to test cultures. *Escherichia coli* strains used for cloning were grown in LB broth or on LB agar. Where noted, tetracycline was used at 5 μg/ml for *Z. mobilis* and at 15 μg/ml for *E. coli*, and kanamycin was used at 100 μg/ml for *Z. mobilis. Z. mobilis* was grown at 30°C and *E. coli* was grown at 37°C.

**Table 1.**
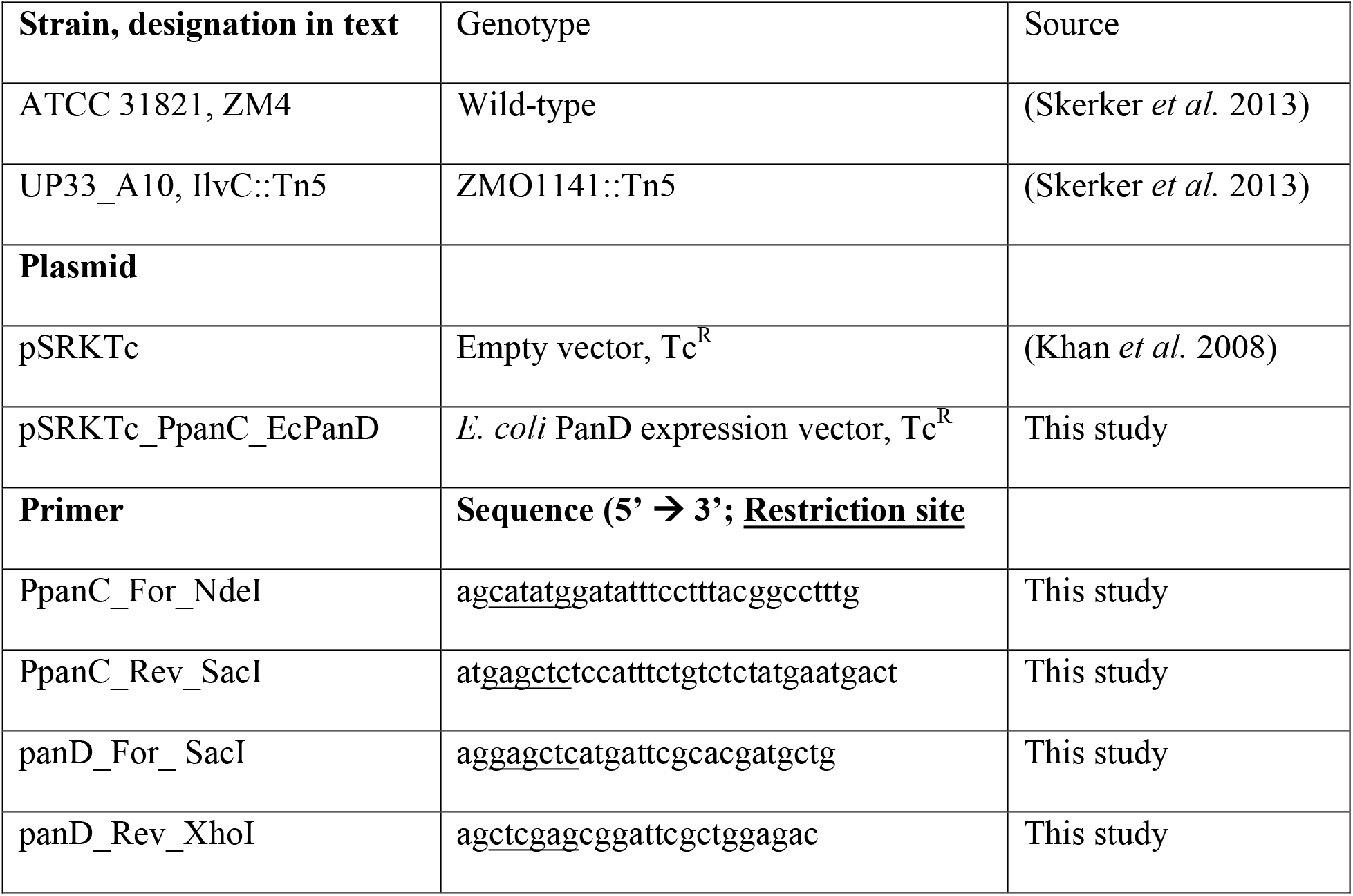
Strains, plasmids, and primers.

### Construction of *Z. mobilis* gene expression vectors

All plasmids and primers are described in Table 1. All enzymes and competent cells were used according to the manufacturer’s instructions. To express *E. coli* PanD in *Z. mobilis*, the ZM4 *panC* promoter (P*panC*) was first amplified from ZM4 genomic DNA using primers to introduce NdeI and SacI restriction sites upstream and downstream of the promoter, respectively. The PCR product was digested with NdeI and SacI (NEB, Ipswich, MA) and then ligated into pSRKTc (Khan *et al*. 2008) that had been digested with the same enzymes. The ligation reaction was used to transform chemically competent *E. coli* NEB10β (NEB) and then cells were plated on selective media. Transformants were screened for pSRKTc with the *PpanC* insert using colony PCR and the correct sequence was confirmed by Sanger sequencing. Next, the *E. coli panD* gene was amplified from *E. coli* MG1655 genomic DNA using primers to introduce SacI and XhoI restriction sites upstream and downstream of the gene, respectively. The gene was then inserted into the pSRKTc_PpanC plasmid downstream of the *PpanC* promoter using the same procedure as above.

All plasmids were transformed into ZM4 by electroporation. ZM4 was first made electro-competent by growing cells to mid-exponential phase in 100 ml YPG, harvesting by centrifugation, washing three times in 10 ml of 10% ice-cold glycerol, and resuspending in 1 ml 10% ice-cold glycerol. Fifty μl aliquots were frozen in an ethanol-dry ice bath and then stored at -80°C. Electroporation was carried out with a BioRad MicroPulse electroporator (Hercules, CA), using program ‘Ecoli 1’, in 1mm electroporation cuvettes. Electroporated cells were allowed to recover in 5 ml YPG for at least 18 hours at 30°C without shaking before plating onto selective media.

### Analytical techniques

Cell densities were monitored by optical density at 660nm (OD_660_) as described (Gordon and McKinlay 2014). Glucose and ethanol were quantified using a Shimadzu (Kyoto, Japan) high performance liquid chromatograph as described (McKinlay, Zeikus, and Vieille 2005).

## Results

### Comparison of pantothenate synthesis operons in *Z. mobilis* isolates

Pantothenate auxotrophy is a common attribute among *Z. mobilis* isolates (Belaich and Senez 1965; De Ley and Swings 1976; Nipkow, Beyeler and Fiechter 1984; Galani, Drainas and Typas 1985; Lawford and Stevnsborg 1986; Cross and Clausen 1993). In fact, a survey of 38 *Zymomonas* sp. isolates found that all strains required pantothenate (De Ley and Swings 1976). In ZM1, a strain for which no genome sequence is publically available, heterologous expression of both *panD* and *panE* eliminated the auxotrophy (Tao, Tomb and Viitanen 2014). To gauge if a lack of *panD* and *panE* might similarly explain the pantothenate auxotrophy in other *Z. mobilis* isolates, we used BLAST (Altschul *et al*. 1990) to look for protein sequences similar to *E. coli* MG1655 PanB, PanC, PanD, and PanE in the seven sequenced *Z. mobilis* strains (Seo *et al*. 2005; Kouvelis *et al*. 2009, 2011; Pappas *et al*. 2011; Desiniotis *et al*. 2012). None of the seven strains had homologs of *panD* or *panE*. Five of the seven strains have putative *panB* and *panC* genes in a region of conserved gene arrangement, or synteny (Fig. 2). One of these five strains, ZM4, had an additional *panBC* gene pair (ZMO1952 and ZMO1971, respectively) 22.4 kb away from the *panBC* pair in this region (ZMO1970 and ZMO1954, respectively), but otherwise there was no syntenty between the two regions. The other two strains, ATCC 29191 and ATCC 29192, lacked genes with any significant sequence similarity to *panB* and *panC* but otherwise showed synteny in this region (Fig. 2).

### *Z. mobilis* ZM4 is auxotrophic for β-alanine but not for pantoate

The absence of *panD* and *panE* suggested that *Z. mobilis* strains having *panB* and *panC* should be incapable of making both β-alanine and pantoate (Fig. 1). In examining the pantothenate synthesis pathway in the metabolism database MetaCyc (Caspi *et al*. 2014), we noticed that other bacteria can make pantoate using IlvC, an enzyme better known for acetohydroxy acid isomeroreductase activity in branched-chain amino acid synthesis. In fact, IlvC and PanE have redundant pantoate synthesis activity in *E. coli* (Elischewski, Pühler and Kalinowski 1999) and IlvC is the only enzyme responsible for pantoate synthesis in *Corynebacterium glutamicum* (Merkamm *et al*. 2003). All seven *Z. mobilis* genomes encode a protein with 37% identity to the *E. coli* IlvC. If *Z. mobilis* can make pantoate using IlvC then the pantothenate auxotrophy could be due to an inability to synthesize β-alanine alone. We therefore tested whether *Z. mobilis* could grow when supplied with β-alanine in place of pantothenate. Growth trends were similar when either β-alanine or pantothenate were provided (Fig. 3A), whereas no growth was observed when both supplements were omitted (Fig. 3A). Providing β-alanine in place of pantothenate also had no effect on the ethanol yield (Fig. 3B).

**Fig 3.**
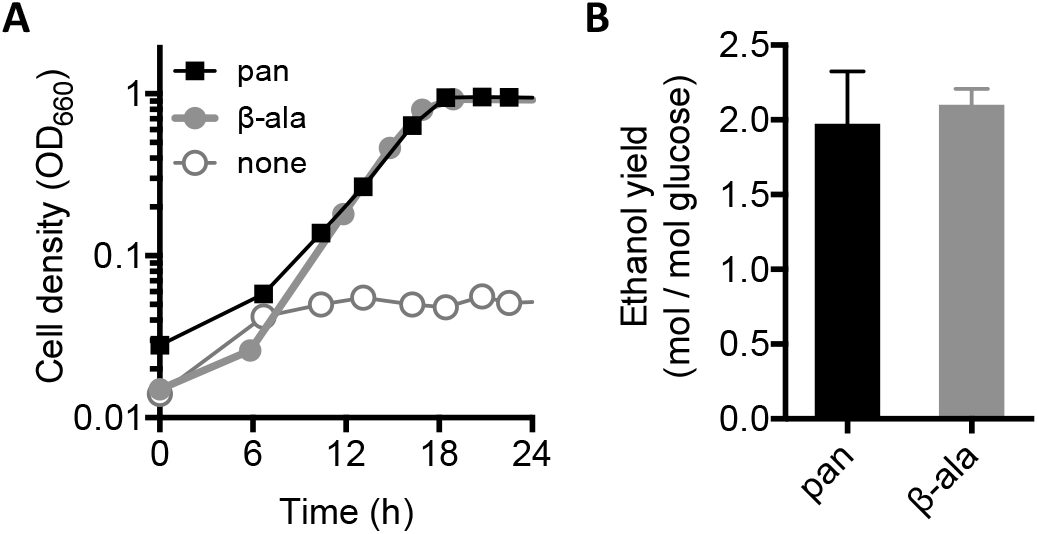
β-alanine (β-ala) can substitute for pantothenate (pan) to support ZM4 growth. **(A)** Representative growth curves in a chemically-defined medium with the specified supplement. Similar trends were observed in at least three replicate cultures. **(B)** Ethanol yields from culture conditions used in panel A. Error bars = SD; n=3.

Since β-alanine can substitute for pantothenate as an essential growth supplement, we hypothesized that the auxotrophy could be eliminated by expressing PanD, the enzyme that produces β-alanine by decarboxylating aspartate (Fig. 1). To test this hypothesis, we constructed a plasmid for expressing the *E. coli panD* gene under control of the *Z. mobilis panC* promoter. This plasmid allowed ZM4 to grow in a medium lacking both pantothenate and β-alanine, whereas an empty vector did not (Fig. 4). We conclude that the *Z. mobilis* ZM4 pantothenate auxotrophy is due to a lack of β-alanine-producing aspartate decarboxylase (PanD) activity and, by extension, that ZM4 has the native capacity to synthesize pantoate.

**Fig. 4.**
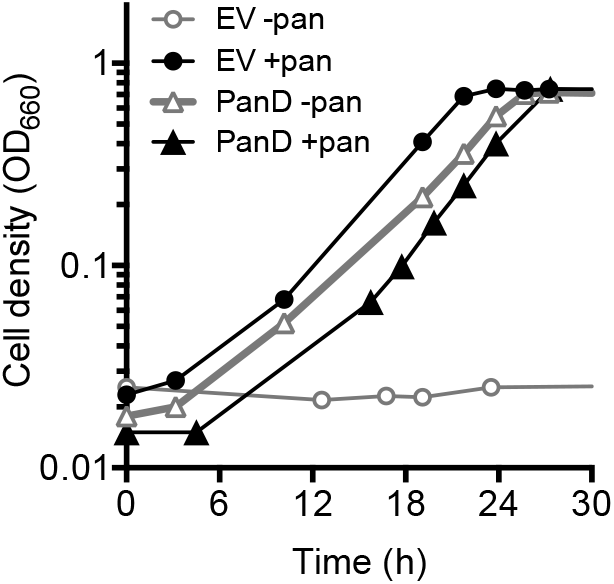
PanD activity allows ZM4 to grow without pantothenate. Representative ZM4 growth curves in a chemically-defined medium with the specified supplement. Similar trends were observed in at least three replicate cultures. EV, empty vector (pSRKTc); PanD, PanD expression vector (pSRKTc_PpanC_EcPanD); pan, pantothenate. All cultures contained tetracycline.

### *Z. mobilis* ZM4 IlvC is responsible for pantoate synthesis

As noted above, IlvC, an enzyme involved in branched-chain amino acid synthesis, can substitute for the pantoate synthesis activity of PanE in other bacteria (Elischewski, Pühler and Kalinowski 1999; Merkamm *et al*. 2003). The ZM4 IlvC gene, ZMO1141, is located 872.6 kb away from the *panBC* pair shown in Figure 2. To test whether IlvC is responsible for pantoate synthesis in ZM4, we examined the growth requirements of an IlvC transposon mutant for pantothenate and β-alanine. The IlvC::Tn mutant could only grow when both pantothenate and branched-chain amino acids were provided (Fig. 5). Unlike WT ZM4, β-alanine in place of pantothenate did not support IlvC::Tn mutant growth. The strict requirement for both pantothenate and branched-chain amino acids by the IlvC::Tn mutant indicates that IlvC is required for *de novo* synthesis of both of these compounds in *Z. mobilis*. (Fig. 5).

**Fig 5.**
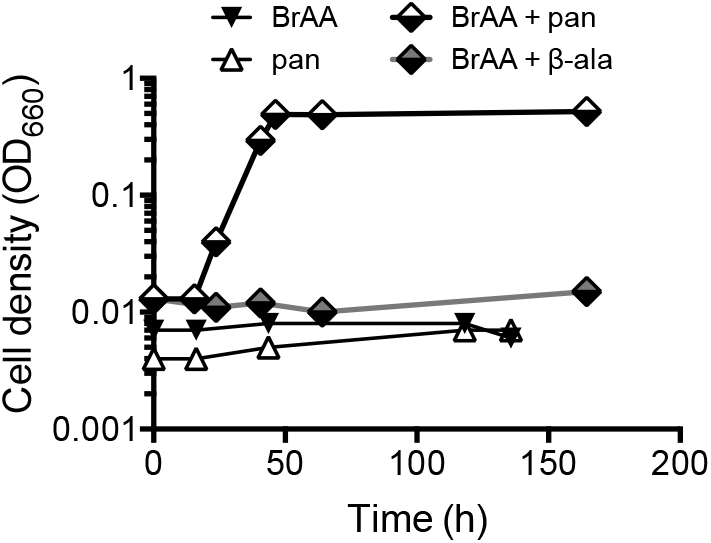
IlvC is required for synthesis of both branched-chain amino acids and pantothenate in ZM4. Representative growth curves for the IlvC::Tn mutant in a chemically-defined medium with the specified supplement. Similar trends were observed in at least three replicate cultures. BrAA, branched-chain amino acids (isoleucine, leucine, and valine); pan, pantothenate; β-ala, β-alanine.

### Discussion

We have demonstrated that the pantothenate auxotrophy in *Z. mobilis* ZM4 is due to the absence of PanD, encoding aspartate decarboxylase (Fig 4). The absence of *panE* does not factor into the auxotrophy, as its absence is compensated for by the activity of IlvC (Fig 5), similar to what has been observed in some other bacteria (Elischewski, Pühler and Kalinowski 1999; Merkamm *et al*. 2003). A patent previously reported that expression of heterologous *panD* and *panE* in ZM1 eliminated the pantothenate auxotrophy; however, the effects of expressing each gene individually was not tested (Tao, Tomb and Viitanen 2014). While the ZM1 genome sequence is not publically available, microarray analysis has shown it to be highly similar to the ZM4 genome sequence (Seo *et al*. 2005). ZM1 is missing 54 genes that are present in ZM4, including the possible second *panB* copy (ZMO1952), but IlvC (ZMO1141) was not among the list of missing genes in ZM1 (Seo *et al*. 2005). Thus, it might only be necessary to express *panD* in ZM1 to eliminate the pantothenate auxotrophy. The same is likely true for other *Z. mobilis* strains that encode *panB* and *panC* (Fig 2). Separately, we found that the product of PanD activity, β-alanine, could substitute for pantothenate in supporting ZM4 growth in a defined medium (Fig 3). β-alanine costs less than a tenth of that of pantothenate and thus can be viewed as a more cost-effective supplement for *Z. mobilis* defined media.

## Acknowledgements

We are grateful to JM Skerker and AP Arkin for providing transposon mutants and the corresponding parental strain, to B LaSarre for manuscript reading, and the McKinlay lab for valuable discussions. This work was supported by the Indiana University College of Arts and Sciences.

## References

Altschul SF, Gish W, Miller W et al. Basic local alignment search tool. J Mol Biol 1990;215:403–10.

Belaich J-P, Senez JC. Influence of aeration and of pantothenate on growth yields of *Zymomonas mobilis*. J Bacteriol 1965;89:1195–200.

Caspi R, Altman T, Billington R et al. The MetaCyc database of metabolic pathways and enzymes and the BioCyc collection of Pathway/Genome Databases. Nucleic Acids Res 2014;42:D459–71.

Cross JS, Clausen EC. Effects of organic buffers on batch fermentations of *Zymomonas mobilis* in a synthetic and complex medium. Biomass and Bioenergy 1993;4:277–81.

Desiniotis A, Kouvelis VN, Davenport K et al. Complete genome sequence of the ethanol-producing *Zymomonas mobilis* subsp. *mobilis* centrotype ATCC 29191. J Bacteriol 2012;194:5966–7.

Elischewski F, Pühler A, Kalinowski J. Pantothenate production in *Escherichia coli* K12 by enhanced expression of the *panE* gene encoding ketopantoate reductase. J Biotechnol 1999;75:135–46.

Galani I, Drainas C, Typas NA. Growth requirements and the establishment of a chemically defined minimal medium in *Zymomonas mobilis*. Biotechnol Lett 1985;7:673–8.

Gordon GC, McKinlay JB. Calvin cycle mutants of photoheterotrophic purple non-sulfur bacteria fail to grow due to an electron imbalance rather than toxic metabolite accumulation. J Bacteriol 2014;196:1231–7.

Jeffries T. Ethanol fermentation on the move. Nat Biotechnol 2005;23:40–1.

Khan SR, Gaines J, Roop RM et al. Broad-host-range expression vectors with tightly regulated promoters and their use to examine the influence of TraR and TraM expression on Ti plasmid quorum sensing. Appl Environ Microbiol 2008;74:5053–62.

Kouvelis VN, Davenport KW, Brettin TS et al. Genome sequence of the ethanol-producing *Zymomonas mobilis* subsp. pomaceae lectotype strain ATCC 29192. J Bacteriol 2011;193:5049–50.

Kouvelis VN, Saunders E, Brettin TS et al. Complete genome sequence of the ethanol producer *Zymomonas mobilis* NCIMB 11163. J Bacteriol 2009; 191:7140–1.

Kremer TA, LaSarre B, Posto AL et al. N_2_ gas is an effective fertilizer for bioethanol production by *Zymomonas mobilis*. Proc Natl Acad Sci 2015;112:2222–6.

Lawford HG, Rousseau JD. Corn steep liquor as a cost-effective nutrition adjunct in high-performance *Zymomonas* ethanol fermentations. Appl Biochem Biotechnol 1997;63-65:287–304.

Lawford HG, Stevnsborg N. Pantothenate limitation does not induce uncoupled growth of *Zymomonas* in chemostat culture. Biotechnol Lett 1986;8:345–50.

De Ley J, Swings J. Phenotypic description, numerical analysis, and proposal for an improved taxonomy and nomenclature of the genus *Zymomonas* Kluyver and van Niel 1936. Int J Syst Bacteriol 1976;26:146–57.

McKinlay JB, Zeikus JG, Vieille C. Insights into *Actinobacillus succinogenes* fermentative metabolism in a chemically defined growth medium. Appl Environ Microbiol 2005;71:6651–6.

Merkamm M, Chassagnole C, Lindley ND et al. Ketopantoate reductase activity is only encoded by *ilvC* in *Corynebacterium glutamicum*. J Biotechnol 2003;104:253–60.

Nipkow A, Beyeler W, Fiechter A. An improved synthetic medium for continuous cultivation of *Zymomonas mobilis*. Appl Microbiol Biotechnol 1984;19:237–40.

Pappas KM, Kouvelis VN, Saunders E et al. Genome sequence of the ethanol-producing *Zymomonas mobilis* subsp. mobilis Lectotype Strain ATCC 10988. J Bacteriol 2011;193:5051–2.

Seo J-S, Chong H, Park HS et al. The genome sequence of the ethanologenic bacterium *Zymomonas mobilis* ZM4. Nat Biotechnol 2005;23:63–8.

Skerker JM, Leon D, Price MN et al. Dissecting a complex chemical stress: chemogenomic profiling of plant hydrolysates. Mol Syst Biol 2013;9:674.

Tao L, Tomb J-F, Viitanen P V. Pantothenic acid biosynthesis in *Zymomonas*. 2014. Patent US8765426 B2.

